# The La Crosse virus M segment determines virus isolate cell-to-cell spread and virulence

**DOI:** 10.64898/2026.09.17.752448

**Authors:** Molly V. Durawa, Nicole C. Rondeau, Sophie N. Spector, Sara A. Thannickal, Abbie E. Weight, Ariana Dedvukaj, Matthew C. Lutchko, Helen M. Lazear, Kenneth A. Stapleford

## Abstract

La Crosse virus (LACV) is an orthobunyavirus spread by mosquitoes in North America and can cause severe neurological disease. Despite this burden, there is a lack of LACV antiviral treatments as our fundamental understanding of how LACV spreads and causes disease remains incomplete. To investigate LACV biology, we took advantage of two genetically similar LACV lineage I isolates (LACV_1960_ and LACV_1978_) where we found that LACV_1960_ infects and replicates at a higher rate than LACV_1978_, while LACV_1978_ exhibits enhanced cell-cell spread and *in vivo* virulence and dissemination. To investigate the genomic determinants behind these phenotypes, we generated reassortants between each isolate. We found that each genomic segment contributed to LACV pathogenesis with the M segment as a dominant determinant for LACV pathogenesis *in vivo* and plaque size and replication *in vitro*. To address which M segment protein contributes to plaque size and infectivity, we generated M segment chimeric viruses using a LACV_1978_ background and swapping in regions of the LACV_1960_ M segment. We found that the Gc head domain determined plaque size and infectivity, with the LACV_1960_ Gc head chimera producing small plaques but having increased infectivity over the wild-type LACV_1978_. Finally, using natural LACV lineage isolates, we showed that plaque size is variable across and within lineages suggesting changes in the M segment may impact LACV in nature. In future studies, we will continue to investigate the mechanisms behind how cell-to-cell spread influences dissemination to better understand how LACV genome segments affect spread, infectivity, virulence, and viral fitness.

**Importance:** Orthobunyaviruses are significant global public health threats. However, our understanding of how these viruses spread and cause disease is not well-defined. In this study, we took advantage of two genetically similar La Crosse virus (LACV) strains that differ in their infectivity, cell-to-cell spread, and virulence. We used reassortant and chimeric LACVs to identify the M segment as a major virulence determinant that regulates cell-to-cell spread and infectivity through the Gc head domain. These studies highlight that different LACV strains behave differently and that the genomic M segment can regulate these phenotypes. Future studies to better dissect how the M segment impacts dissemination will provide important avenues for antiviral development.

## Introduction

The *Orthobunyavirus* genus within the *Peribunyaviridae* family is comprised of arthropod-borne viruses that cause disease in both humans and livestock (1). These viruses, which include Oropouche virus, La Crosse virus (LACV), and Jamestown Canyon virus, among others, can result in febrile and neuroinvasive disease (2–4). However, there is a dearth of treatment or prevention options beyond avoiding bites by their arthropod vectors. For this reason, it is critical to better understand the mechanisms of orthobunyavirus pathogenesis to generate viable therapeutics.

LACV, a mosquito-borne orthobunyavirus, is the leading cause of arboviral pediatric encephalitis in North America, yet our understanding of how LACV spreads and causes disease is incomplete. LACV consists of three genetically distinct lineages, which generally separate geographically (5–7). Different strains of LACV, even within the same lineage, have been found to exhibit different levels of replication and virulence (8, 9), suggesting that viral genetic determinants contribute to disease. LACV is a tri-segmented, negative-sense RNA virus with a genome consisting of the S, M, and L segments. The S segment encodes the nucleocapsid protein N and interferon antagonist nonstructural protein NSs (10–12), and the L segment encodes the RNA-dependent RNA polymerase (RdRp) (13, 14). The M segment encodes the surface glycoproteins Gn and Gc needed for virion assembly and infection (15–18) as well as the protein NSm required for assembly and transmission(19). While previous work has defined the roles of each protein, much remains unknown about how these determinants regulate the virulence and spread of LACV and other orthobunyaviruses.

In these studies, we sought to better understand the contribution of LACV genomic segments by comparing the spread and virulence of genetically similar LACV strains. To do this, we used two LACV lineage I isolates for which we have infectious clone systems, LACV_1960_ and LACV_1978_. We studied the cell-to-cell spread, infectivity, binding, replication, and *in vivo* virulence and dissemination of these two wild-type viruses as well as of segment reassortant viruses and M segment chimeras to understand the genetic basis of phenotypic differences. We found that the LACV_1978_ isolate had enhanced cell-to-cell spread, virulence, and neuroinvasiveness while the LACV_1960_ isolate had enhanced infectivity and replication *in vitro.* Moreover, we determined that the M segment and more specifically, the Gc head domain. Moreover, we found that LACV virulence was driven by multiple genomic segments through inter-segment interactions. These findings will inform further research into the determinants of pathogenesis in LACV and other orthobunyaviruses.

## Results

### Two non-natural mutations in the LACV_1960_ infectious clone S segment attenuates LACV replication and virulence

Previous studies have shown that different La Crosse virus (LACV) strains exhibited differences in viral growth kinetics and pathogenesis (8, 9). However, the molecular determinants driving these phenotypes are not well-defined. To begin characterizing these viral determinants, we acquired infectious clones of the lineage I isolates, LACV_Human/1960_ (LACV_1960_) (20) and the LACV_Mosquito/1978_ (LACV_1978_) (21). We first looked to compare the genetic similarities between these LACV isolates at the protein and RNA level (**Figure 1A**, **Tables 1 and 2**). At the protein level, we found that the S segment proteins N and NSs were 100% conserved while there were 30 and 17 amino acid differences between M and L segment proteins, respectively (**Table 1**). At the RNA level, the S, M, and L segments were variable between strains, with the M segment being the least conserved at 95.7% nucleotide conservation (**Table 2**).

**Figure 1:**
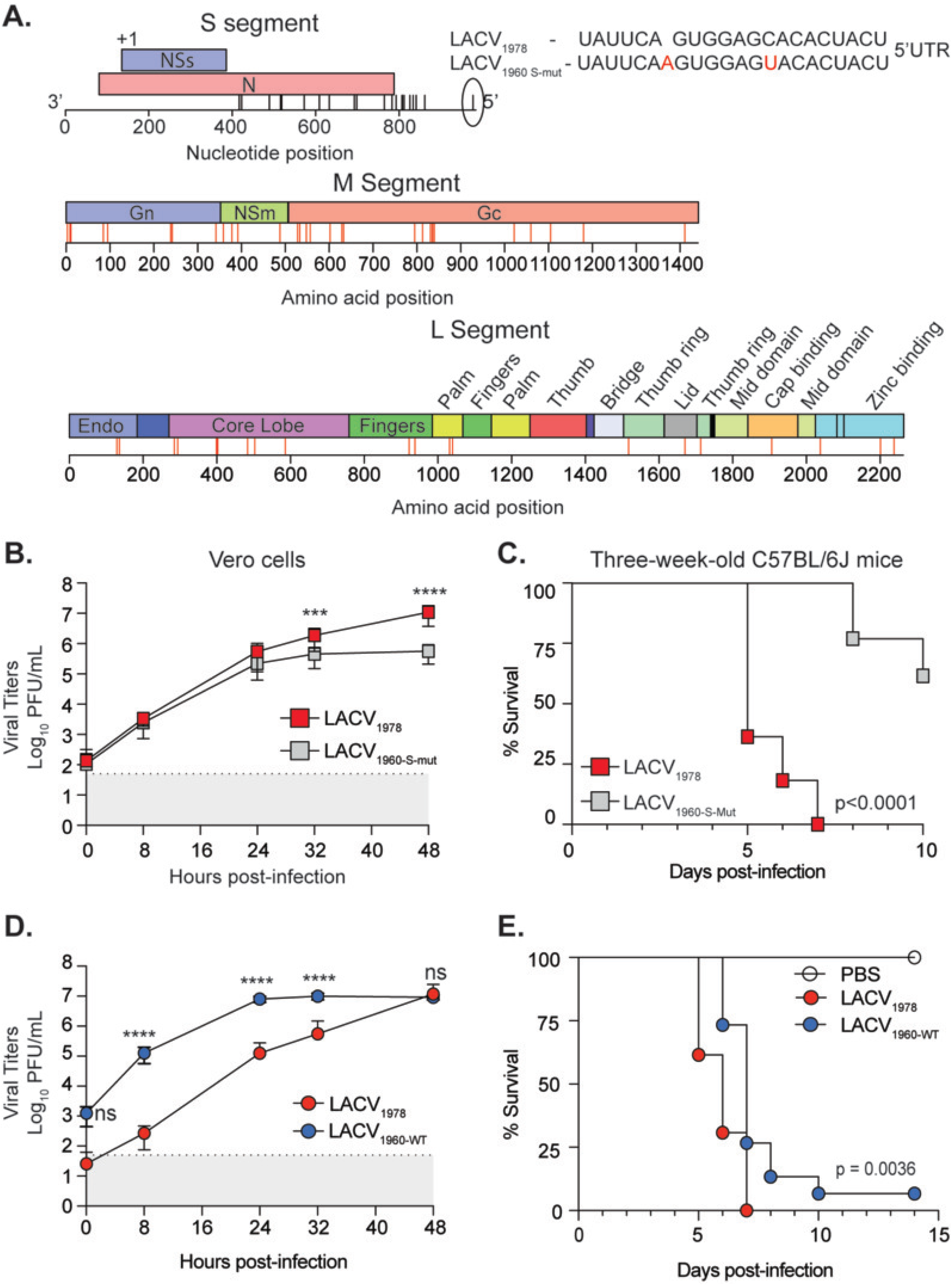
Two non-natural mutations in the LACV 1960 infectious clone attenuate replication and virulence. **A.** Left: RNA (S segment) and protein (M and L segment) alignments of LACV_1960_ and LACV_1978_ infectious clones. Black ticks indicate nucleotide differences. Red ticks indicate amino acid differences. Right: S segment inset showing S segment 5’ UTR sequences. Variant nucleotides are in red. **B.** Vero cells were infected with WT LACV 1978 and the 1960 S-mut virus at a MOI of 0.1 The supernatant was collected and viral titers quantified by plaque assay. Data represent the mean and standard deviation (SD) of three independent experiments with internal duplicates, N=6. Mann-Whitney test. *** p<0.001, **** p<0.0001. **C.** Three-week-old C57BL/6J mice were inoculated via the left rear footpad with 50,000 PFU of WT LACV_1978_ or the LACV_1960_ S-mut virus. Mice were monitored and weighed daily for 10 days. Data represent five independent infections (WT LACV_1978_ N=13; LACV_1960_ S-mut N=19). Mantel-Cox test. **D.** Vero cells were infected with WT LACV_1978_ and WT LACV_1960_ at a MOI of 0.1 Viral titers in the supernatant were quantified by plaque assay. Data represent the mean and SD of five independent experiments with internal duplicates, N=8-10. Mann-Whitney test. ns = not significant. **** p<0.0001 **E.** Three-week-old C57BL/6J mice were inoculated as in C. with WT LACV 1978 and WT LACV 1960. Mice were monitored and weighed daily for 14 days. Data represent three independent experiments. (WT LACV_1978_ N=13; WT LACV_1960_ N=15). Mantel-Cox test.

**Table 1:** Protein conservation between LACV strains.

|  | LACV<sub>Human/1960</sub><br>S segment | LACV<sub>Human/1960</sub><br>M segment | LACV<sub>Human/1960</sub><br>L segment |
| --- | --- | --- | --- |
| LACV<sub>Mosquito/1978</sub><br>S segment | 0 AA<br>100% |  |  |
| LACV<sub>Mosquito/1978</sub><br>M segment |  | 30 AA<br>97.9% |  |
| LACV<sub>Mosquito/1978</sub><br>L segment |  |  | 18 AA<br>99.2% |

*Aligned using ClustalW Multiple Sequence alignment Number of amino acid differences indicated above consensus percentage*

**Table 2:** RNA conservation between LACV strains.

|  | LACV<sub>Human/1960</sub><br>S segment | LACV<sub>Human/1960</sub><br>M segment | LACV<sub>Human/1960</sub><br>L segment |
| --- | --- | --- | --- |
| LACV<sub>Mosquito/1978</sub><br>S segment | 22 nt<br>97.8% |  |  |
| LACV<sub>Mosquito/1978</sub><br>M segment |  | 195 nt<br>95.7% |  |
| LACV<sub>Mosquito/1978</sub><br>L segment |  |  | 284 nt<br>95.9% |

*Aligned using Smith-Waterman RNA alignment*

Interestingly, we observed two nucleotide changes in the LACV_1960_ infectious clone S segment 5’ UTR (20) that were not found in nature (**Figure 1A, right**). These changes were a C to U mutation and an A insertion in the infectious clone S segment sequence. Given the role of the 5’UTR in orthobunyavirus replication and transcription (22–25), we hypothesized that these changes would influence virus replication *in vitro* and *in vivo*. To test whether the LACV_1960_ with the two mutations was attenuated compared to the LACV_1978_ isolate, we infected Vero cells with the WT LACV_1978_ virus and the 1960 S segment mutant (1960-S-mut) at an MOI of 0.1, collected supernatant at 0, 8, 24, 32, and 48 hours post-infection (hpi), and quantified viral titers by plaque assay. While the 1960-S-mutant replicated at similar levels to WT LACV_1978_ for the first 24 hours, the S-mut virus was significantly attenuated later during infection (**Figure. 1B**). To address virulence of the LACV_1960_ S-mut virus, we infected three-week-old C57BL/6J mice with 50,000 PFU of each virus via the footpad and monitored survival over the course of 10 days (**Figure 1C**). For the WT LACV_1978_ virus, all mice succumb to infection by seven days post-infection. However, the LACV_1960_ S-mutant was significantly attenuated in mice with only 25% mice succumbing to infection by day 10. To test how these two nucleotides impact replication, we then generated a new LACV 1960 S segment infectious clone plasmid lacking these two nucleotide changes and repeated these *in vitro* and *in vivo* experiments (**Figure. 1D and E**). We found that in contrast to the 1960-S-mutant, the WT LACV_1960_ virus replicates to significantly higher titers in Vero cells compared to WT LACV_1978_ (**Figure 1D**). Moreover, we are now able to restore virulence of the WT LACV_1960_ virus in three-week-old C57BL/6J mice (**Figure 1E**). These results show that the two non-natural S segment mutations attenuate the LACV infectious clone. We used the new WT LACV_1960_ system for all subsequent studies.

### LACV_1978_ has enhanced cell-to-cell spread

While generating viral stocks we observed that LACV_1978_ generated significantly larger plaques than LACV_1960_ (**Figure 2A**). This phenotype was also observed with the 1960-S-mut virus. Because plaque assays allow the movement of virus only between adjacent cells, we predicted that differences in cell-to-cell spread are responsible for the difference in plaque size between LACV_1960_ and LACV_1978_. To test this hypothesis, we used immunofluorescence to measure the size of viral foci, allowing us to quantify cell infection rather than cell death. We infected Vero cells with each virus and stained with LACV anti-sera at 16, 24, and 48 hours post-infection (**Figure 2B and D**). We observed that LACV_1978_ foci grew more rapidly than those of LACV_1960_, a trend that was recapitulated in human myoblast cells, ruling out that this phenotype was unique to Vero cells (**Figure 2C and E**). These results suggest that some LACV isolates may use cell-to-cell spread more efficiently than other isolates.

**Figure 2:**
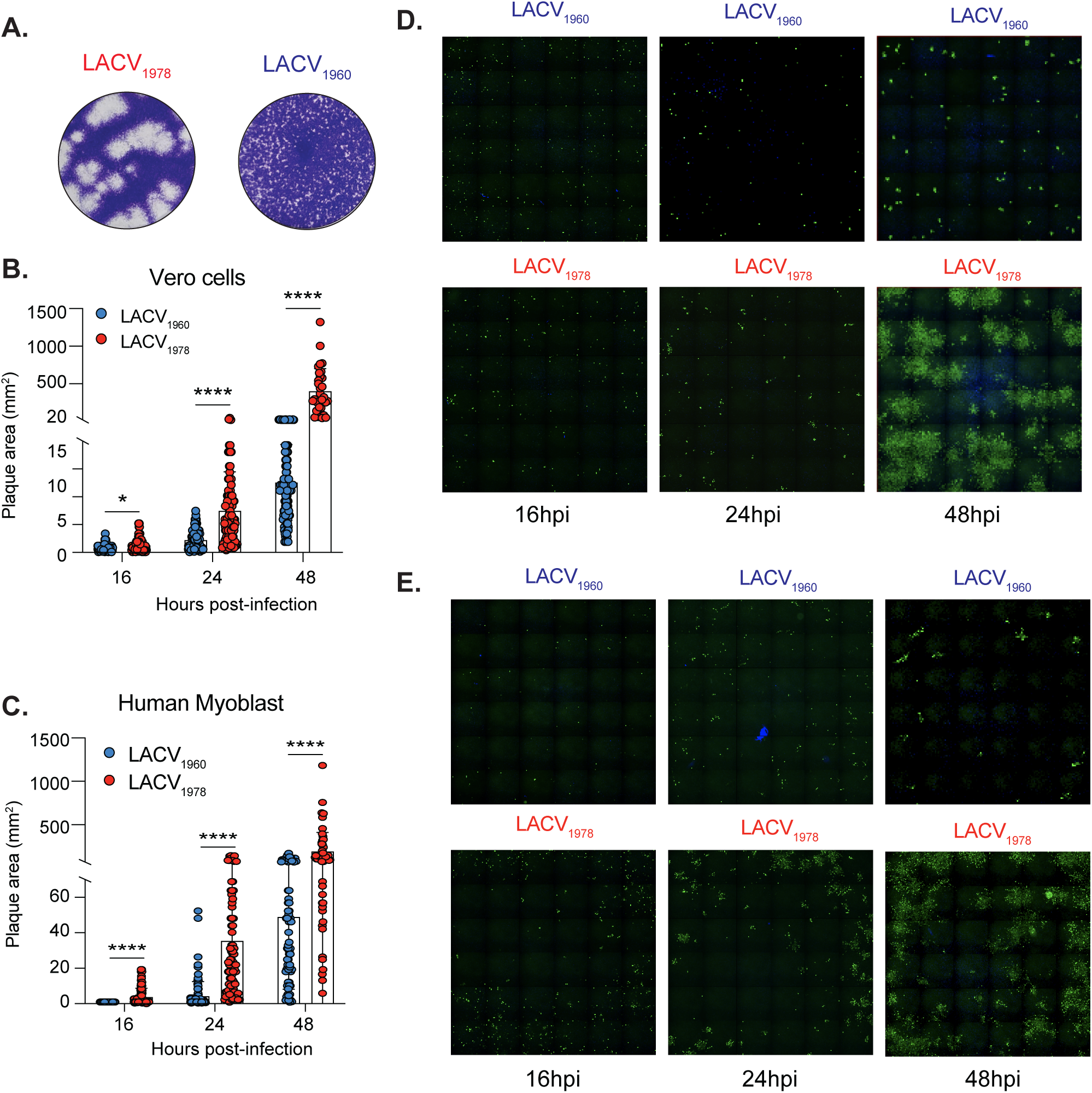
LACV_1978_ has enhanced cell-to-cell spread. **A.** Representative plaque images from WT LACV_1978_ and WT LACV_1960_ on Vero cells at three days post incubation. **B and D.** Vero cells were infected with 10-fold serial dilutions of each virus for 1 hour followed by the addition of an agarose overlay. At 16, 24, and 48 hours, cells were fixed and stained with anti-LACV antisera and anti-rabbit Alexa488 secondary antibody. The plaque area was quantified using Image J. **C and E.** Human myoblasts were infected and processed as in B and D. Data represent the mean and SD from a total of 26-100 plaques quantified from two independent experiments. Mann-Whitney test. * p<0.05, **** p<0.0001.

### LACV_1960_ has increased replication in Aag2 cells and enhanced infectivity in Vero cells

Given the enhanced replication of the WT LACV_1960_ isolate in Vero cells, we then asked whether this phenotype was similar for other cell types. We infected human myoblasts, important cells that serve as the initial site of LACV infection(26), at an MOI of 0.1, collected supernatant at 0, 8, 24, 32, and 48 hours post-infection, and used plaque assays to quantify viral titers (**Figure 3A**). While we did observe an increase in replication of LACV_1960_ over LACV_1978_ in myoblasts, these results were not statistically significant. However, when we infected *Aedes aegypti* Aag2 cells at an MOI of 0.1 and collected time points at 0, 8, and 24 hours post-infection, we found that the LACV_1960_ virus grew to higher titers in mosquito cells (**Figure 3B**).

**Figure 3:**
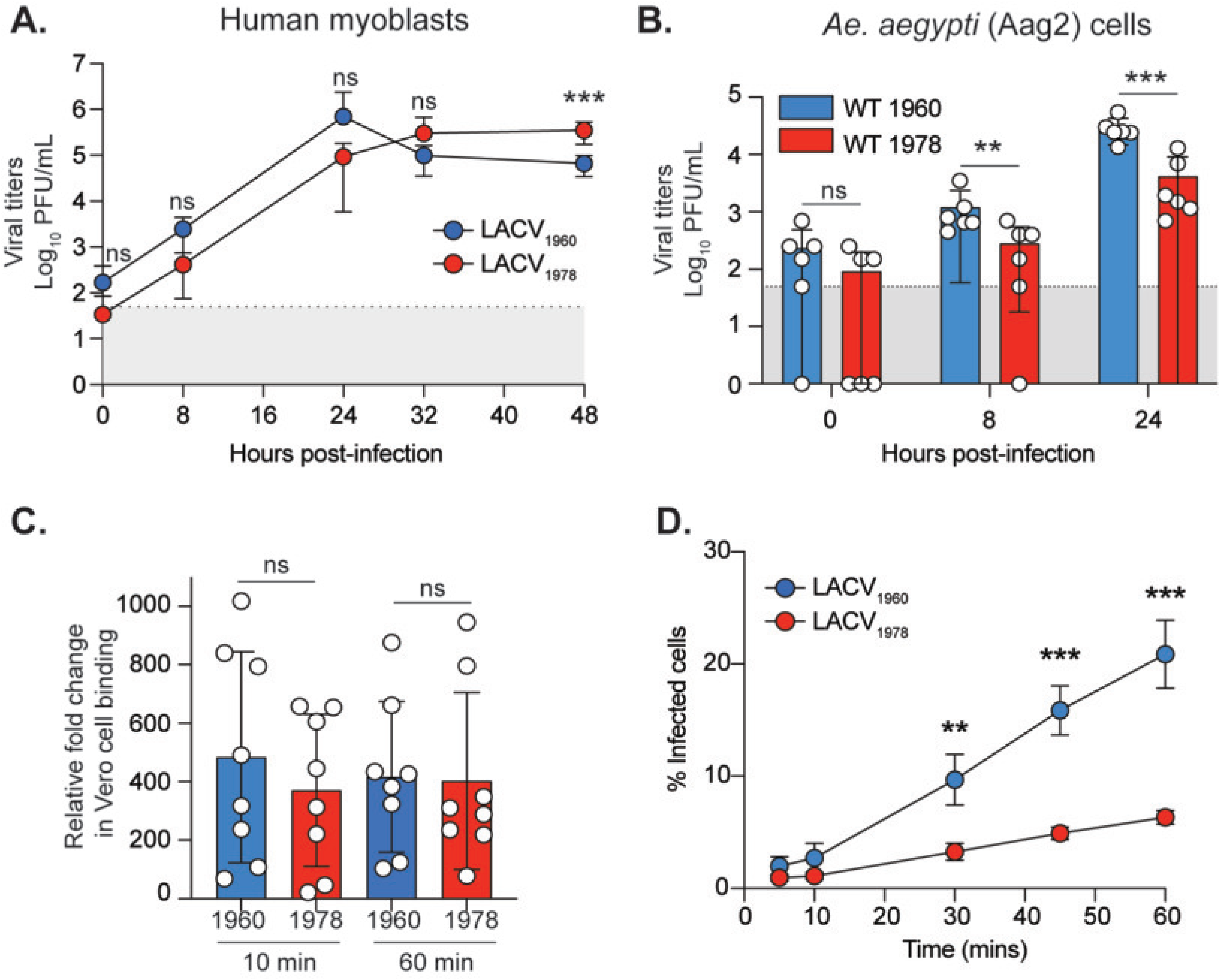
LACV1960 has enhanced replication in Aag2 cells and enhanced infectivity in Vero cells. **A.** Human myoblasts were infected with each virus at a MOI of 0.1. At each time point, the supernatant was collected and infectious virus quantified by plaque assay. Data represent the mean and SD of four independent experiments with internal duplicates N=6-8. Mann-Whitney test. *** p<0.001. **B.** *Ae. Aegypti* Aag2 cells were infected with each virus at a MOI of 0.1 and analyzed as in A. Three independent experiments. N=6. Mann-Whitney t-test. ** p<0.01, *** p<0.001. **C.** Vero cell binding assay. Vero cells were incubated for one hour at 4°C in the presence of 20 mM NH_4_Cl to block entry. Cells were then incubated with each gradient purified virus at a MOI of 100 based on RNA genomes. After 10 and 60 min of incubation, the cells were washed three times with cold PBS and RNA extracted with Trizol. S segment and GAPDH RNA was quantified by RT-qPCR and the data represented as 2^-ΔΔCt^. Data represent the mean and SD of four independent experiments with internal duplicates N=8. Mann-Whitney test. All values are not significant. **D.** Virus infectivity assays. Vero cells were incubated with each virus at a MOI of 1. At 5, 10, 30, 45, and 60 mins post-infection, 20 mM NH4Cl was added to the culture to block further virus spread. 24 hours post-infection, cells were fixed and stained with anti-LACV antigen and DAPI. The number of infected cells was quantified using the CellInsight CX7 high-content microscope. Data represent the mean and SD of three independent experiments with internal duplicates N=6. Mann-Whitney test. ** p<0.01, *** p<0.001.

We next sought to determine the mechanism for the increase in replication of LACV_1960_ compared to LACV_1978_ in Vero cells despite the reduced cell-to-cell spread. We first hypothesized that LACV_1960_ may have improved cell binding, leading to more infected cells. To investigate this hypothesis, we performed binding assays by first treating Vero cells with 20 mM ammonium chloride (NH_4_Cl) for one hour on ice to block endocytosis. We then added sucrose-purified virus (MOI=100 based on RNA genomes) on ice for 10 or 60 minutes, washed the cells, extracted RNA, and quantified viral S segment RNA by RT-qPCR. We found no significant difference in binding between LACV_1960_ and LACV_1978_ at either timepoint (**Figure 3C**). We next investigated if there was a difference in the rate of infectivity between the two viruses. We infected Vero cells with each virus at an MOI of 1 and then blocked further infection with 20 mM NH_4_Cl at 5, 15, 30, 45, and 60 minutes post-infection. We found that LACV_1960_ exhibited significantly higher infection at 30, 45, and 60 minutes post-infection, with more than triple the proportion of infected cells compared to LACV_1978_ at 60 mins post-infection (**Figure 3D**). These results suggest that LACV_1960_ has greater infectivity compared to LACV_1978_, leading to more infected cells and enhanced replication even in the absence of robust cell-to-cell spread.

### LACV_1978_ has increased replication *in vivo*

Since LACV_1978_ has enhanced virulence and enhanced cell-to-cell spread *in vitro*, we then asked whether LACV_1978_ exhibited increased replication in mice compared to LACV_1960_. To characterize the replication phenotypes of the two strains *in vivo*, we inoculated three-week-old C57BL/6-J mice with 50,000 PFU of LACV_1960_ or LACV_1978_ via left hind footpad injection. We harvested the footpad (site of inoculation) and brain at three days post-infection and quantified infectious virus and viral RNA in each tissue (**Figure 4**). Mice infected with LACV_1978_ had higher overall titers in the footpad compared to mice infected with LACV_1960_, with average LACV_1978_ titers having a more than ten-fold increase over LACV_1960_ (**Figure 4A**). Furthermore, there were three LACV_1960_ infected mice with no virus detected in the footpad, whereas the footpads of all LACV_1978_-infected mice contained detectable virus (**Figure 4A**). LACV_1978_ additionally appeared to disseminate to the brain faster, with nearly all LACV_1978_ infected mice having virus detected in the brain compared to only one LACV_1960_-infected mouse (**Figure 4A**). These results—obtained using plaque assay— were confirmed via RNA quantification of the same samples using RT-qPCR (**Figure 4B**) and show that while the majority of mice have LACV_1978_ RNA in the brain at three days post-infection, LACV_1960_ was largely absent confirming that virus had not yet reached the brain by day 3 post-infection.

**Figure 4:**
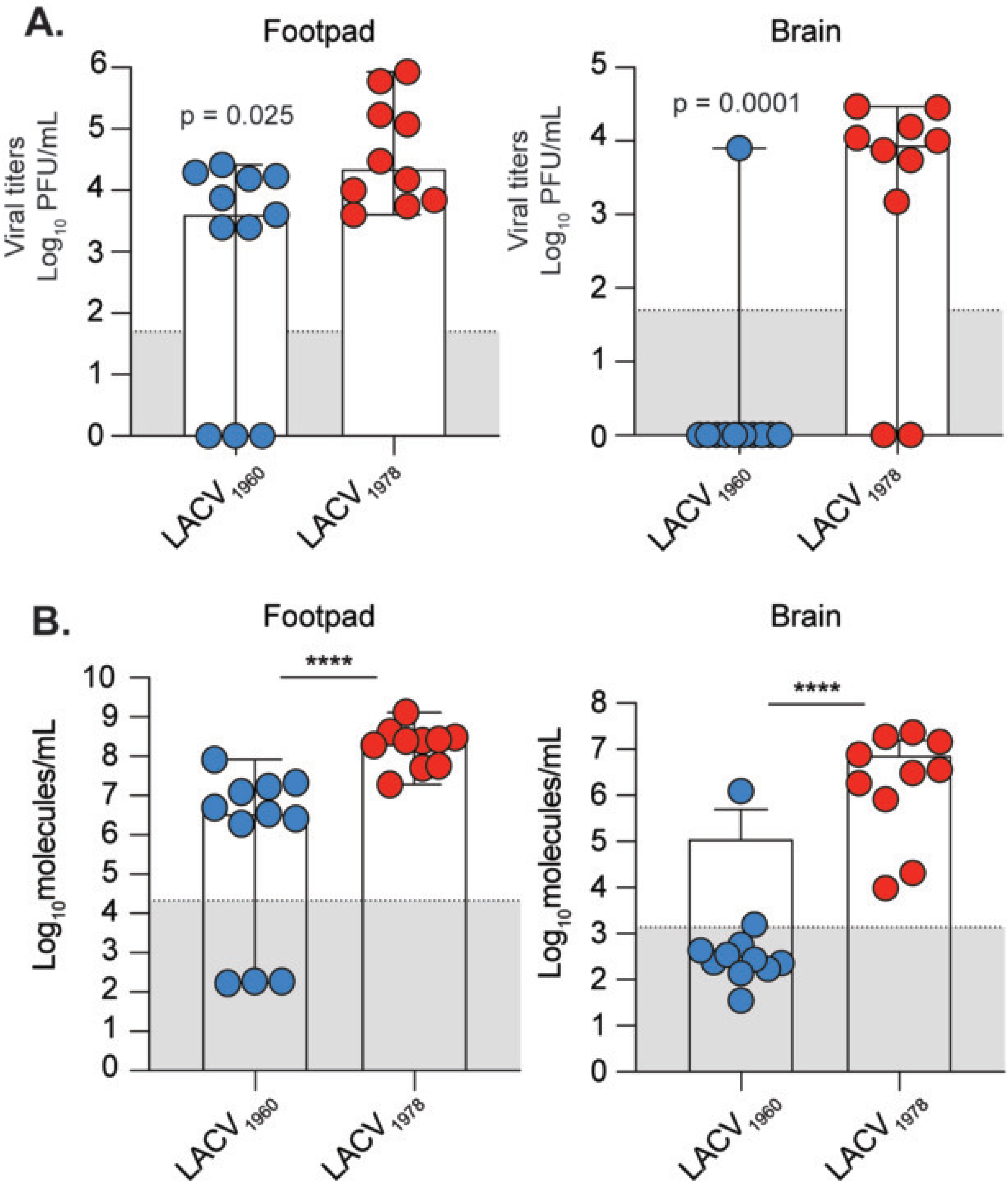
LACV_1978_ reaches the brain faster than LACV_1960_ Three-week-old C57BL/6J mice were inoculated with 50,000 PFU of each virus or PBS control in the left rear footpad. At three days post-infection, mice were euthanized and the footpad (site of inoculation) and brain were harvested to quantify (**A**) viral titers by plaque assay and (**B**) LACV S segment RNA by RT-qPCR. Data represent the median and range of two independent experiments. LACV_1978_ N=10; LACV_1960_ N=11. Mann-Whitney test. P values are shown in the figure. **** p < 0.0001.

### The LACV M segment contributes to plaque size and virulence

Given that we had found that LACV_1978_ spread faster cell-to-cell and exhibited greater virulence, whereas LACV_1960_ had enhanced replication and infectivity but lower virulence, we asked which part of the viral genome dictated these phenotypes. The proteins encoded by the orthobunyavirus M segment have been shown to be responsible for attachment, entry, and cell-to-cell spread, so we hypothesized that the LACV M segment would likewise be responsible for the observed differences between LACV_1960_ and LACV_1978_. To test this hypothesis, we used our infectious clone system to generate reassortant viruses by transfecting baby hamster kidney (BHK) BSR T7/5 cells with different combinations of LACV_1960_ and LACV_1978_ S, M, and L plasmids (**Figure 5**). When we generated each reassortant, we first observed that only the M segment led to a complete reversal of the plaque phenotype (**Figure 5A**) suggesting that the M segment is responsible for cell-to-cell spread.

**Figure 5:**
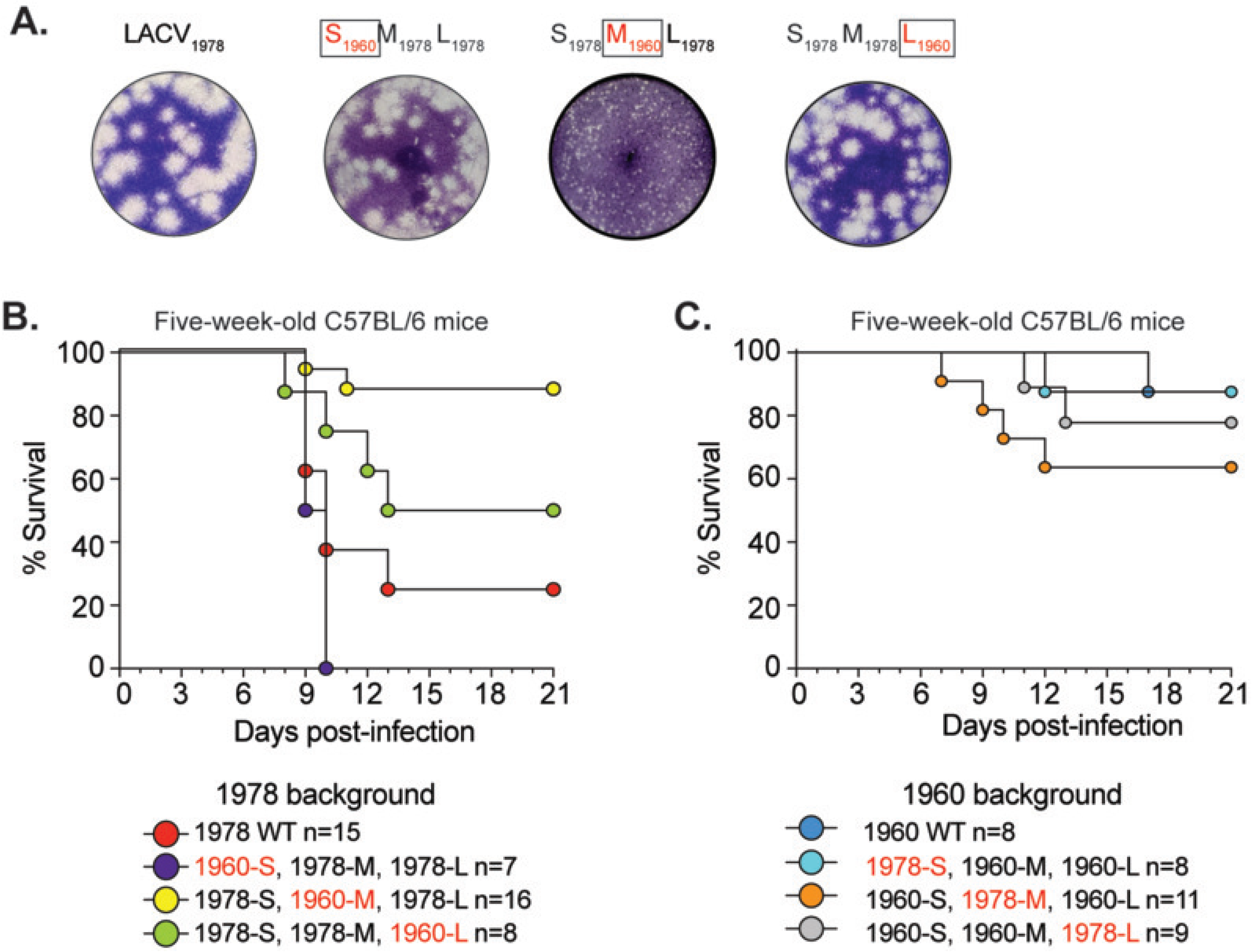
LACV M segment regulates plaque size and is a major virulence determinant. **A.** Representative plaque images of WT LACV_1978_ and the reassortants containing the LACV_1960_ S, M, and L segments on Vero cells at three days post incubation. **B and C.** Five-week-old C57BL/6 mice were inoculated with 10,000 PFU of each virus by subcutaneous footpad injection. Mice were monitored for disease signs and humane endpoints for 21 days.

We then used a five-week-old mouse model to test how each LACV segment contributes to virulence. The five-week-old mouse model was used to better distinguish virulence phenotypes compared to the more susceptible three-week-old mouse model. We found that the LACV_1978_ virus was more virulent than LACV_1960_ with 70% of mice succumbing to LACV_1978_ compared to only ∼10% succumbing to LACV_1960_ (**Figure 5B-C**, consistent with the relative virulence of these strains in three-week-old mice (**Figure 1**). When we introduced the 1960 S, M, and L segments into the 1978 background, we found that the S segment increased virulence (100% of mice succumbing) whereas the L segment reduced virulence slightly (∼50% of mice succumbing). However, the 1960 M segment had the greatest impact on virulence with only ∼10% of mice succumbing to infection, similar to WT LACV_1960_ (**Figure 5B**). Using the reassortants on the 1960 background, we found that interestingly the LACV_1978_ S and L segments had no impact on virulence, whereas the 1978 M segment was able to increase virulence slightly (∼40% of mice succuming) (**Figure 5C**). These results indicate the M segment is a major virulence determinant.

### The LACV M segment influences replication *in vitro* and dissemination in mice

Given the enhanced replication of the LACV_1960_ virus in Vero and Aag2 cells, we then asked whether changing the M segment to the 1978 would reduce replication *in vitro*. We infected Vero cells (**Figure 6A**), human myoblasts (**Figure 6B**), and Aag2 cells (**Figure 6C**) with each WT virus and the 1978 M reassortant at a MOI of 0.1. We collected supernatants at various time points and quantified viral titers by plaque assay. In all cell types tested, we found that the 1978 M segment on the 1960 background reduced replication *in vitro*.

**Figure 6:**
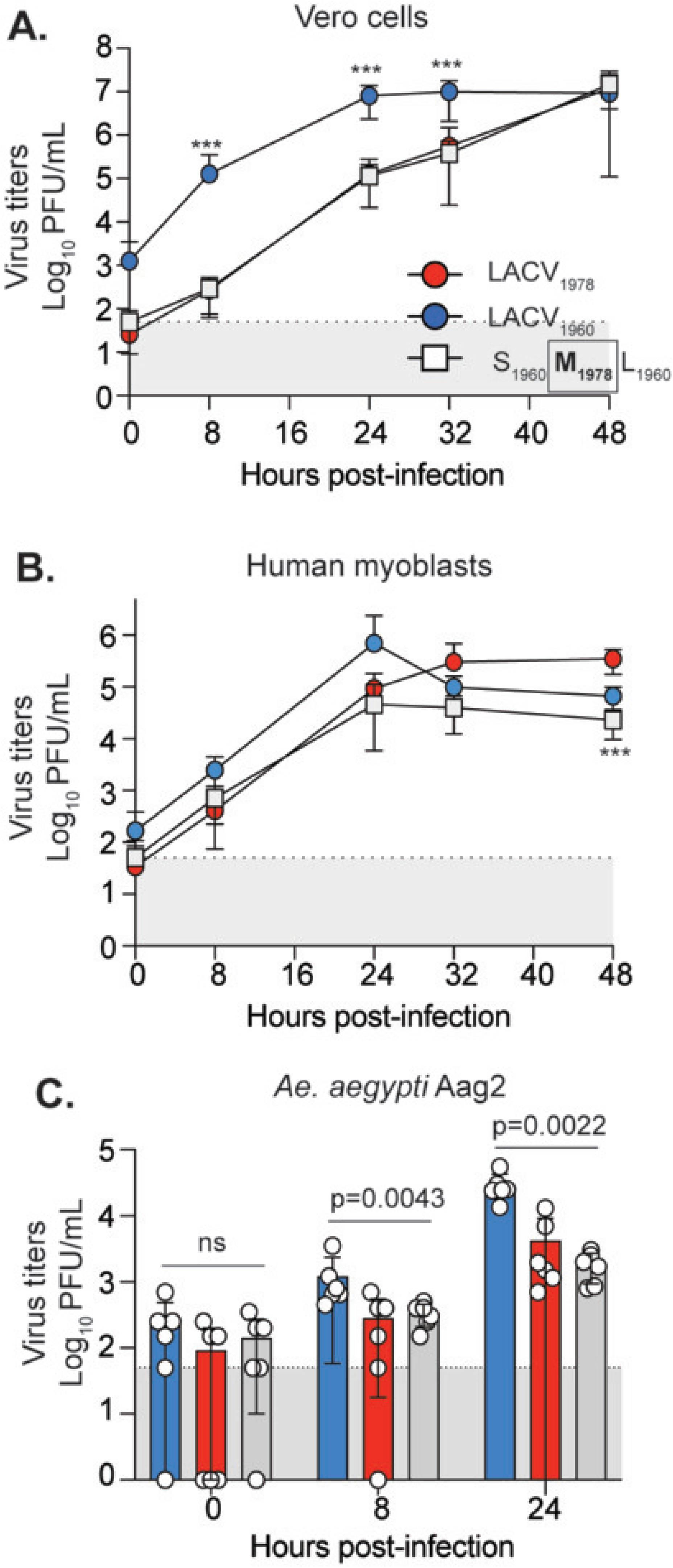
The LACV_1978_ M segment reduces replication *in vitro*. Vero cells (**A**), Human myoblasts (**B**), or Aag2 cells (**C**) were infected with each virus at a MOI of 0.1 At each time point, the supernatant was collected and viral titers were quantified by plaque assay. Data represent the mean and SD of three independent experiments for each cell line with internal duplicates N=6. Kruskal-Wallis test. P values are shown. ns = not significant.

We next asked whether the M segment contributed strain-specific effects on LACV replication in mice, comparing the 1960 M and 1978 M segments on the 1978 background. We inoculated three-week-old C57BL/6J mice via left hind footpad injection with 50,000 PFU of LACV_1978_ or the S_1978_M_1960_L_1978_ reassortant. All LACV_1978_-inoculated mice succumbed to infection by seven days post-infection, whereas only approximately 50% of S_1978_M_1960_L_1978_-inoculated mice had at the same timepoint (**Figure 7A**). We then measured viral replication at days three and five post-infection. In the footpad, the two viruses replicated to similar titers (**Figure 7C and D**). However, while the viral titers in the brain at five days post-infection were similar, at three days post-infection, only one mouse inoculated with S_1978_M_1960_L_1978_ had detectable virus in the brain compared to over half of the mice inoculated with LACV_1978_ (**Figure 7C**). Together, these results show that the 1978 M segment can significantly influence replication *in vitro* and *in vivo*.

**Figure 7:**
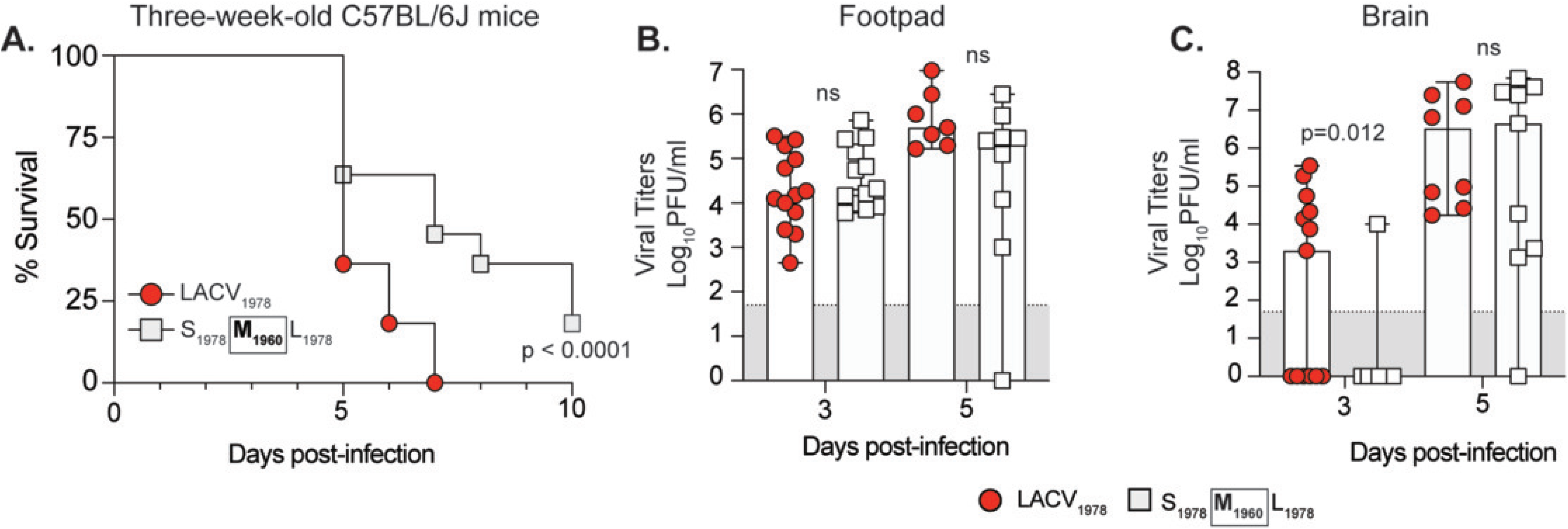
The LACV_1960_ M segment attenuates virulence and dissemination to the brain. **A.** Three-week-old C57BL/6J mice were inoculated with 50,000 PFU of WT LACV_1978_ or the M_1960_ reassortant via the left rear footpad. Mice were monitored and weighed daily for 10 days. Data represent three independent experiments. (LACV_1978_ N=11; M_1960_ reassortant N=11). Mantel-Cox test. **B.** Mice were inoculated as in A. Mice were euthanized at three and five dpi and the footpad and brain harvested. Viral titers were quantified in the footpad and brain by plaque assay. Data represent the median and range of two to three independent experiments. Day 3 LACV_1978_ N=13; M_1960_ reassortant N=13. Day 5 LACV_1978_ N=8; M_1960_ reassortant N=9). Mann-Whitney test. ns = not significant.

### The LACV Gc head domain contributes to plaque size and infectivity

To determine which M segment proteins influences differences in infectivity and spread between LACV strains, we generated chimeric viruses in which specific proteins of the LACV_1978_ M segment were swapped with LACV_1960_ (**Figure 8A**). Consistent with previous observations, the WT LACV_1978_ virus produced large plaques, and the M swap (S_1978_M_1960_L_1978_) produced small plaques (**Figure 8B**). The Gn and NSm swap viruses produced plaques similar to those of WT LACV_1978_, but when we replaced the region encoding Gc or the Gc head domain with that of LACV_1960_, the plaques presented the smaller phenotype of WT LACV_1960_ (**Figure 8B**). We then characterized the infectivity of the chimeric viruses. Relative to WT LACV_1978_, only the full M swap and Gc head domain swaps had significantly higher infectivity, both with an average infectivity of more than double that of WT LACV_1978_ (**Figure 8C**). Interestingly, the full Gc swap did not recapitulate the full M swap, suggesting that the combination of Gn, NSm, and Gc play important roles in LACV infectivity. These studies highlight the role of the Gc head domain in driving cell-to-cell spread and infectivity. Finally, we have previously shown that the LACV Gc head domain is variable between LACV lineages(17), allowing us to hypothesize that different LACV lineages may have different plaque sizes. To address plaque size between LACV lineages we performed plaque assays using 12 LACV strains from lineage I (5 viruses), lineage II (4 virus), and lineage III (4 viruses) (**Figure 9**). Interestingly, there was a large variance in plaque size, even between viruses of the same lineage. Together, these results indicate that differences between LACV isolate M segments may dictate LACV cell-to-cell spread.

**Figure 8:**
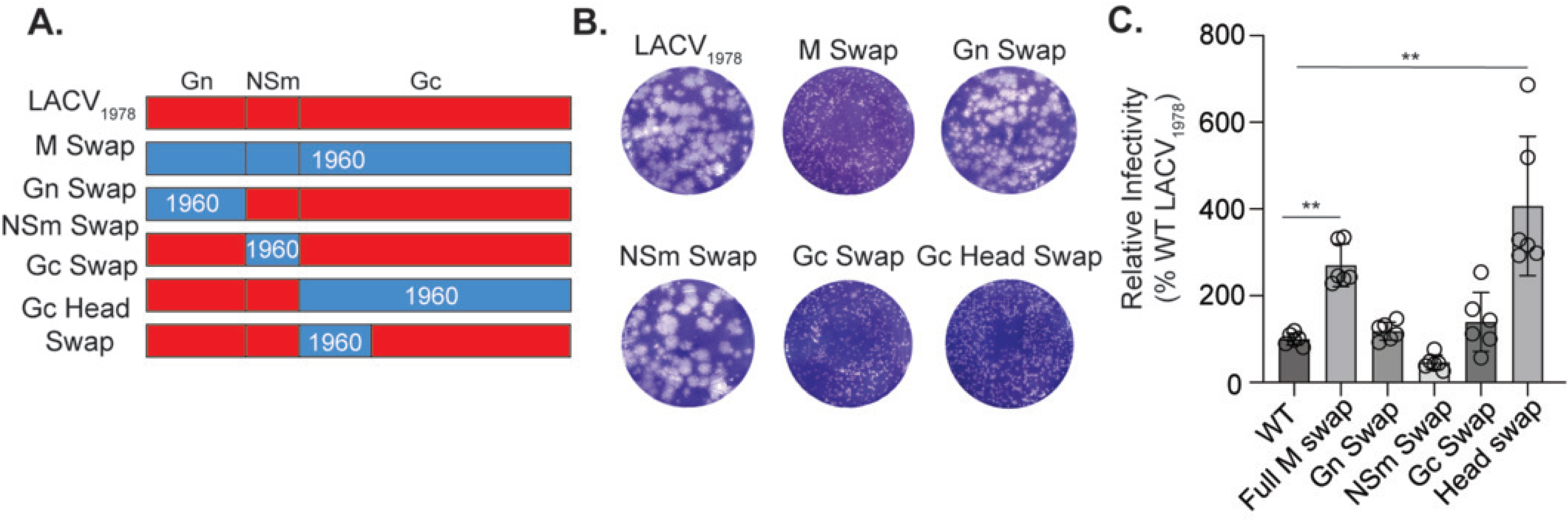
The LACV M segment head domain contributes to plaque size and infectivity. **A.** Schematic of LACV M segment chimera reassortants. **B.** Representative plaque assay images of each virus on Vero cells at three days post incubation. **C.** Virus infectivity assay. Vero cells were infected with each virus at an MOI of 1 for 60 min followed by the addition of 20 mM NH_4_Cl. 24 hpi, cells were fixed and stained with anti-LACV anti-sera and DAPI. The number of infected cells was quantified using the high-content microscope. Data represent the mean and SD of three independent experiments with internal duplicates. N=6. Kruskal-Wallis test. ** p<0.01.

**Figure 9:**
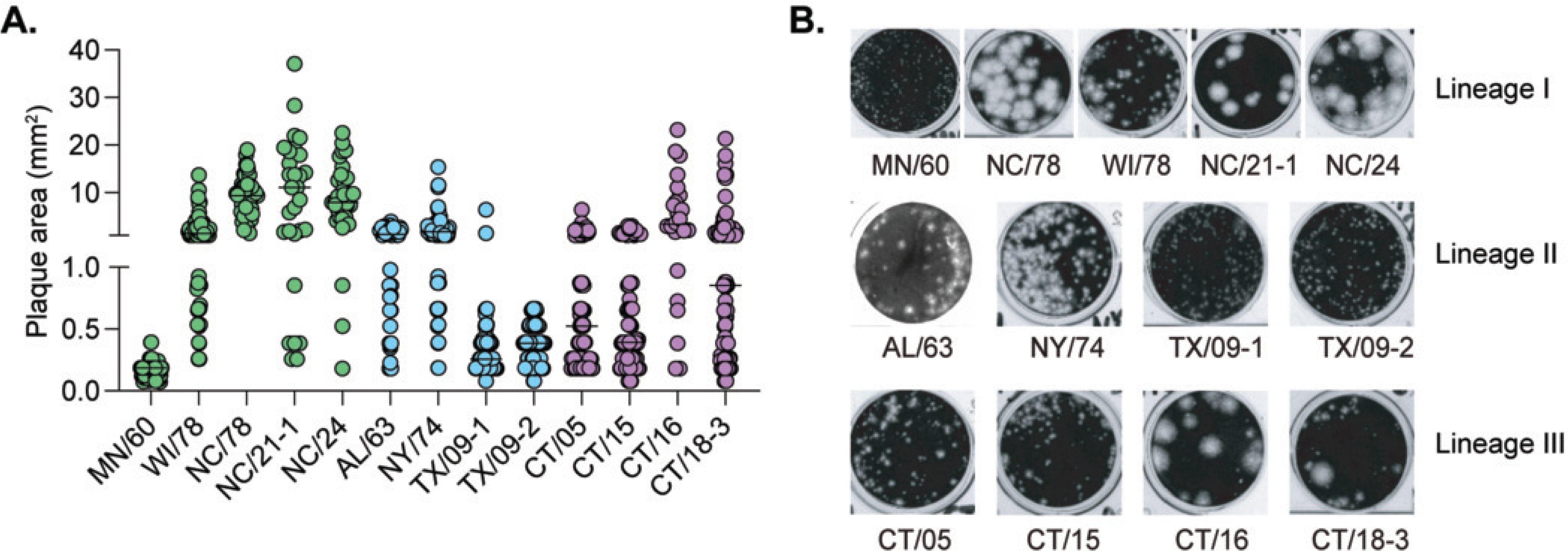
LACV isolates display varying plaque sizes. **A.** LACV isolates were plaqued on Vero cells, developed three days post incubation, and plaque size was quantified by Image J. 24-116 plaques were quantified from two independent infections. **B.** Representative plaque assay images developed at three days post infection.

## Discussion

Despite the public health burden imposed by orthobunyaviruses, there are no antiviral therapies or treatments. To better protect against these viruses, it is critical to understand how they establish disease and the mechanisms that determine virulence. Previous orthobunyavirus studies have taken advantage of their segmented genomes to generate reassortant viruses to study the impacts of genomic segments on pathogenicity (2, 27). However, these studies utilize reassortants composed of different viral species, which differ significantly at the genome level. Therefore, we aimed to investigate virulence determinants between two genetically similar LACV lineage I strains, LACV_1960_ and LACV_1978_.

We found that the two genetically-similar viruses had very different plaque phenotypes, with LACV_1960_ producing small plaques and LACV_1978_ producing large plaques—which we determined to be due to enhanced cell-to-cell spread by LACV_1978_. While large plaque size might be expected to correspond to enhanced viral replication, we found that unexpectedly LACV_1960_ replicated better in Vero cells and mosquito Aag2 cells compared to LACV_1978_ as well as having enhanced infectivity. These results suggest that different LACV isolates have evolved to use cell-to-cell spread at the expense of enhanced infectivity to drive spread. Better understanding host responses to LACV infection, including in neurons and other relevant cell types, will be important for defining strain-specific pathogenic mechanisms.

We found that LACV_1978_ was more virulent than LACV_1960_ with faster replication within the initial site of inoculation and dissemination to the brain. Our experiments with reassortant viruses revealed that the M segment drives the differences we observed in plaque phenotype, replication, and virulence, similar to what has been seen for other orthobunyaviruses (27–29). When we swapped the S, M, and L segments of LACV_1960_ and LACV_1978_, the plaque size and replication kinetics were determined by the M segment; i.e., viruses with the LACV_1978_ M segment produced larger plaques and had attenuated replication, and viruses with the LACV_1960_ M segment produced smaller plaques and had enhanced replication. The M segment was similarly responsible for virulence in mice and the rate of viral dissemination to the brain. The enhanced virulence of viruses with the LACV_1978_ M segment may indicate a critical role for cell-to-cell spread in dissemination and neuroinvasion, as viruses with the LACV_1978_ M segment were more virulent and produced greater viral loads in the brain despite their reduced *in* infectivity and replication in cell culture. LACV neuroinvasion can occur through breakdown of the blood brain barrier as a result of vascular leakage in the olfactory bulb(30), further study into the specific role of the M segment proteins in facilitating this process will be important in helping to identify therapeutic targets.

We found that the Gc head domain appears to be responsible for the differences in plaque size and infectivity between LACV_1960_ and LACV_1978_. The Gc head domain, which forms the tip of the Gc class II glycoprotein spike, has been shown to play a critical role in LACV replication, infectivity, assembly, and virulence and dissemination(15, 17, 31, 32) consistent with our findings. Our results, in conjunction with previous studies that have found Gc and the head domain specifically to be targets of neutralizing antibodies, can help explain differences in pathogenicity between LACV strains (or, indeed, other orthobunyaviruses) and to discover therapeutics (32–34).

Additionally, our observation that a full Gc swap had no significant impact on infectivity— but did result in a change in plaque size—is interesting. It is possible that the physical spike structure is affected by the interactions of the different Gc domains with each other or with the Gn glycoprotein—which forms a heterodimer with the Gc glycoprotein—such that swapping in the LACV_1960_ Gc head domain enhances infectivity but swapping in the entire Gc region does not. The position of the Gc head domain at the tip of the spike suggests a potential role for it in receptor binding—although this is seemingly at odds with our finding that there is no significant difference in binding between LACV_1978_ and LACV_1960_. The role of the Gc head domain in receptor binding, as well as how it specifically impacts infectivity, is an additional avenue for further study.

While we have illustrated here that the M segment and Gc head domain play a crucial role in infectivity and cell-to-cell spread, there are certainly other factors that contribute to LACV pathogenesis. Future studies that interrogate other regions of the M segment (e.g. through point mutations), as well as the S and L segments, and their impacts on cell-to-cell spread, neuroinvasion, and other determinants of virulence will be critical. Additional investigation of the chimeric viruses used in this study—e.g. in mice, in human myoblasts and neuronal cell models— will also help to inform the specific roles played by the Gc head domain and the other regions of the M segment. Going forward, we hope that our findings on the role of the M segment and Gc head domain in cell-to-cell spread and virulence will help to inform further investigation into and treatment of LACV and other orthobunyaviruses.

## Acknowledgements

We thank all members of the Stapleford Lab for helpful discussions on this manuscript. This work was funded by the NYU Grossman School of Medicine Startup (KAS), NIAID/NIH R01 AI62774-01A1 (KAS), R01 AI170625 (HML), R01 AI203471 (HML), T32 AI007419 (AEW), and F31 AI202949 (AEW).

## Methods

### Cell lines

BHK BSR/T7 cells, a gift from Dr. Steven Whitehead at the National Institutes of Health (NIH) (21), were cultured in Dulbecco’s Modified Eagle Medium (DMEM) supplemented with 10% heat-inactivated fetal bovine serum (FBS), 1% non-essential amino acids (NEAA), and 10 mM HEPES. 1 mg/mL geneticin was added in alternate passages to maintain T7 selection. Vero cells (CCL-81, American Type Culture Collection [ATCC]) were cultured in DMEM supplemented with 10% newborn calf serum (NBCS). Human myoblasts, a gift from Dr. Michael Kyba at the University of Minnesota(26), were cultured in HAM’s F10 Nutrient Mixture supplemented with 20% FBS, 1% GlutaMax, 10 ng/mL human basic fibroblast growth factor, 40 ng/mL dexamethasone, and 100 µM beta-mercaptoethanol. Mammalian cells were cultured at 37°C with 5% CO_2_. *Aedes aegypti* Aag2 cells, a gift from Dr. Paul Turner at Yale University(35), were cultured in DMEM supplemented with 10% FBS, 1% NEAA, and 10 mM HEPES at 28°C with 5% CO_2._ All cells were tested monthly for mycoplasma.

### Viruses

Wild-type La Crosse viruses mosquito/1978 (LACV_1978_)(21) and human/1960 (LACV_1960_)(20) were generated using an infectious clone system from Dr. Stephen Whitehead (LACV_1978_) and Dr. Friedmann Weber (LACV_1960_), respectively. The wild-type LACV_1960_ S segment was synthesized at Twist Biosciences and sub-cloned into the original infectious clone plasmid. LACV M segment chimeras were generated via overlapping PCR and standard molecular biology techniques using the primers in **Table 3**. All plasmids were sequenced at Plasmidsaurus. The natural LACV isolates WI/78, NC/21-1, NC/24, AL/63, NY/74, TX/09-1, TX/09-2, and CT/05 were obtained from Dr. Brandy Russell at the Center for Disease Control and Prevention Arbovirus Reference Center and isolates CT/15, CT/16, and CT/18 were obtained from Dr. Philip Amstrong at the Connecticut Agriculture Experimental Station(5, 36). Each isolate was amplified once on Vero cells to generate a working stock and titered by plaque assay.

**Table 3:**
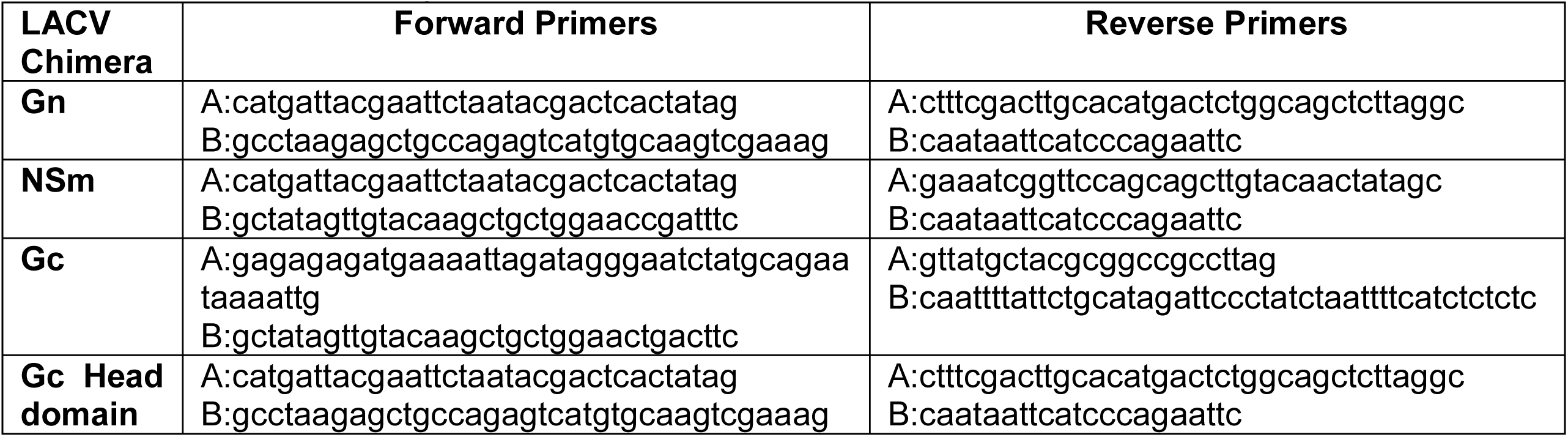
Primers used to generate LACV chimeras.

To generate recombinant virus, BHK BSR/T7 cells (seeded 200,000 cells/well in a 6-well plate 1 d prior) were transfected with 2 µg of plasmid for each genomic segment (S, M, and L) in a mixture containing 250 µL Opti-MEM (Gibco) and 18 µL TransIT-LT1 transfection reagent (Mirus). Cells were incubated for 1 d at 37°C with 5% CO_2_, transfection media was removed, and complete media was added. Cells were incubated and monitored daily for cytopathic effects, at which point the supernatant was collected, centrifuged at 1,200 rpm for 5 mins, aliquoted, and stored at −80°C. Working viral stocks were generated by infecting a monolayer of Vero cells with this transfection product, incubating at 37°C with 5% CO_2_, and monitoring the cells. Upon observation of major cytopathic effects, the supernatant was harvested, centrifuged at 1,200 rpm for 5 mins, aliquoted, and stored at −80°C.

### Growth curves

The day prior to infection, cells were seeded 55,000 cells/well (Vero cells and human myoblasts) or 200,000 cells/well (Aag2 cells) in 24-well plates. Each virus was diluted in DMEM and added to cells at an MOI of 0.1. Following a 1 h incubation at 37°C, cells were washed twice with PBS, and complete media was replaced. For Vero cells and human myoblasts, 100 µL of supernatant was collected at 0, 8, 24, 32, and 48 hours post-infection (hpi), and an equivalent volume of complete media was replaced. For Aag2 cells, the supernatant was collected at 0, 8, and 24 hpi. Plaque assay was used to quantify viral titer.

### Virus spread assay

Each virus was serially diluted 10-fold in DMEM and added to a monolayer of cells in 12-well plates (350,000 Vero cells or human myoblasts/well). Plates were incubated for 1 h at 37°C before the addition of a 0.8% agarose overlay (in DMEM supplemented with 2% NBCS for Vero cells or complete media for human myoblasts). Plates were incubated again for 16, 24, or 48 h and subsequently fixed with 4% formalin for 1 h at room temperature. Cells were stained with LACV anti-sera and DAPI as described below.

### Plaque assay

Viral samples were serially diluted 10-fold in DMEM and added to a monolayer of Vero cells in a 12-well plate (350,000 cells/well). Plates were incubated for 1 h at 37°C, and then 1.5 mL of overlay (DMEM with 0.8% agarose, 2% NBCS, and 1% Antibiotic-Antimycotic for mouse samples) was added to each well. Plates were returned to the incubator for 72–96 h and then fixed with 4% formalin. After 1 h, the agarose overlay was removed, and cells were stained with crystal violet. Plaques were counted at the lowest countable dilution to quantify viral titers.

### Immunostaining

All of the following incubations occurred at room temperature. After fixation, cells were washed twice with Perm/Wash (BD Biosciences), permeabilized with 0.25% TritonX-100 for 10 minutes, and blocked for 1 h with a blocking buffer (0.2% BSA and 0.05% Saponin in PBS). A 1:2,000 dilution of primary rabbit anti-LACV antibodies (a gift from Dr. Karin Peterson at the NIH) in blocking buffer was added for 2 h. Cells were then washed three times with Perm/Wash, and a dilution of secondary goat anti-rabbit IgG Alexa488 (1:2,000) and DAPI (4′,6-diamidino-2-phenylindole (1:1,000)) in blocking buffer was added for 1 h before washing with Perm/Wash three times. PBS was added to all wells, and the number of infected cells was quantified using the CX7 CellInsight high-content microscope.

### Mouse infections

All animal work was approved by the Institutional Animal Care and Use Committee (IACUC) at the NYU Grossman School of Medicine (IA16-01783) and the University of North Carolina at Chapel Hill (Protocol #25-019). Infections completed at the NYU Grossman School of Medicine: Three-week-old wild-type C57BL/6J mice (Strain #000664, Jackson Laboratory – bred in-house) were anesthetized with isofluorane and inoculated with either PBS or 50,000 PFU of each virus diluted in PBS via needle inoculation in the left rear footpad. Mice were then monitored daily for 10 days for signs of disease. Mice exhibiting neurological disease signs or greater than 20% body weight loss were humanely euthanized. In addition, mice infected as described above were euthanized 3 dpi, and brain and footpad samples were harvested in tubes containing 1 mL of plaquing media (DMEM supplemented with 10% NBCS and 1x Antibiotic-Antimycotic) and metal beads. Samples were homogenized for 10 min (footpad) or 5 min (brain) using a Pro Series Bullet Blender homogenizer. Samples were then centrifuged for 3 mins at 10,000 *x g*, and clarified supernatant was used in a plaque assay to quantify viral titer as described. For RNA analysis, 250 μL of clarified supernatant was added to 500 μL of Trizol, and RNA was extracted as described below.

Five-week-old wild-type C57BL/6 mice (bred in-house) were anesthetized using isoflurane and inoculated by subcutaneous footpad injection with 10,000 PFU of each virus diluted in Hank’s balanced salt solution with Ca^2+^ and Mg^2+^ (HBSS) and 1% heat-inactivated FBS. Mice were monitored daily for disease signs for 21 days. Mice that reached humane endpoint, defined as the loss of ≥20% of the starting weight, nonresponsiveness, or severe neurologic disease signs (hunching, seizure, paralysis) that interfered with the ability to access food and water, were humanely euthanized and scored as dead on the following day.

### RNA extractions, cDNA synthesis, and qPCR

RNA was isolated in TRIzol reagent (ThermoFisher Scientific) and extracted using the Direct-zol RNA Miniprep Plus Kit (Zymo Research) according to the manufacturer’s instructions. cDNA was generated using the Maxima H Minus First-Strand cDNA Synthesis Kit (Thermo Fisher Scientific) using random primers and the manufacturer’s instructions. RT-qPCR analysis was completed with Power SYBR Green (Applied Biosciences) and either GAPDH (control) or S segment primers as described(17). For the binding assay, relative fold change in binding compared to an untreated control was calculated using 2^-ΔΔCt^. For quantification of RNA from mouse samples, a standard curve was generated using serial dilutions of the S segment plasmid.

### Virus binding assay

Purified viral stocks were generated via ultracentrifugation of virus supernatant over a 20% sucrose cushsion in PBS at 25,000 rpm for 4 h followed by resuspension of pellet in plaquing media. 100 μL of sample was added to 400 μL of Trizol, and RNA was extracted. Viral RNA was quantified via RT-qPCR. For viral binding assays, Vero cells were seeded in 24-well plates at 55,000 cells/well. Media was removed from cells, and 0.5 mL of a 20 mM ammonium chloride solution in complete media was added to each well. Following a 1 h incubation at 4°C, media was removed, and 100 μL of either plaquing media or virus (MOI = 100 based on viral S segment RNA molecules) was added to each well. Plates were kept on ice for either 10 or 60 min and then washed three times with PBS. 250 uL of Trizol was added to each well for RNA extraction and quantification as described above.

### Virus infectivity assay

Vero cells were seeded 15,000 cells/well in a 96-well plate the day prior to infection. On the day of infection, media was removed, cells were washed with PBS, and cells were incubated with each virus at an MOI of 1 at 37°C. At 5 min, 15 min, 30 min, 45 min, and 60 min post incubation, 20 mM NH_4_Cl in complete media was added to the respective wells to block further infection. Cells were incubated for 24 hours at 37°C and then fixed with 4% paraformaldehyde for 30 m prior to staining as described above.

### Sequence alignments

The LACV_1978_ and LACV_1960_ infectious clone sequences were aligned using SnapGene (v 8.2.2).

### Statistics and data analysis

All data were analyzed using GraphPad Prism (v 11.0.2). All *in vitro* experiments were completed at least three independent times with internal duplicates. All *in vivo* infections were completed at least two independent times. Statistic tests are denoted in the figure legends. P-value <0.05 was considered statistically significant.

## Data availability

All data are present in this manuscript.

